# Construction of an inducible amyloid expression circuit in *Bacillus megaterium:* A case study with CsgA and TasA

**DOI:** 10.1101/858266

**Authors:** Elin M. Larsson, John B. McManus, Richard M. Murray

**Affiliations:** Department of Bioengineering, California Institute of Technology, 1200 East California Blvd, Pasadena, CA, 91125, USA; Army Research Laboratory – West Campus, California Institute of Technology, 1200 East California Blvd, Pasadena, CA, 91125, USA; Control and Dynamical systems, California Institute of Technology, 1200 East California Blvd, Pasadena, CA, 91125, USA

## Abstract

Environmental applications of synthetic biology such as water remediation require engineered strains to function robustly in a fluctuating and potentially hostile environment. The construction of synthetic biofilm formation circuits could potentially alleviate this issue by promoting cell survival. Towards this end, we construct a xylose-inducible system for the expression of the functional amyloids CsgA and TasA in the soil bacterium *Bacillus megaterium*. We find that although both amyloids are expressed, only TasA is successfully exported from the cells. Furthermore, expression of CsgA results in a significant growth penalty for the cells while expression of TasA does not. Finally, we show that TasA expression conveys a small but detectable increase in cells’ adhesion to nickel beads. These results suggest that TasA is a promising candidate for future work on synthetic biofilm formation in *B. megaterium*.

## Introduction

Water contamination by a variety of compounds such as heavy metals is becoming a global health issue [1]. Consequently, the demand for high-performance, low-cost, detection and remediation methods are increasing. Microbial biosensors are an emerging solution to this problem in which synthetic biology is used to construct genetic detection circuits [2]. In the case of environmental remediation, one such function can be the ability to sense the presence of toxins and secrete toxin-degrading compounds in response to detection. However, it is important that the chassis be robust to the environment selected for deployment. Since biofilm formation is a common survival strategy for microbial communities in natural environments [3,4], it is possible that when engineering a community, the robustness of the community is increased when cells are clustered in a biofilm-like state. The ultimate aim is therefore to engineer a sensing and remediating community immobilized at the site of contamination. In this study, we explore expression of amyloid fibers in *B. megaterium* as a potential strategy to promote inducible biofilm formation.

### Biofilm formation

A biofilm is an aggregate composed of different microbial species held together by an extracellular matrix made up of compounds produced by the community members [3]. Being in a biofilm increases the microbial community’s resistance to environmental stress, antibiotics and the immune response of host organisms [4]. The matrix offers protection to the cells and can facilitate the exchange of nutrients within the community [3]. There are many examples of biofilm-forming microbial communities in nature that facilitate environmental remediation. One such example is wastewater treatment plants where microbial communities degrade nutrients and hazardous chemicals. The communities can be made up of planktonic cells or stuck to a particle in the form of a biofilm [5].

Colonization by biofilm formation is a complex process that involves several steps, the first of which is cell adhesion. Following adhesion is cell growth, and finally, cell detachment [3]. The cells can adhere to each other, to abiotic surfaces or to surfaces covered with a conditioning film made up of various organic and inorganic matter [6]. Biofilm formation is facilitated by the production of exopolymeric compounds (EPSs), adhesion proteins and by integral membrane flagella or pili [6]. Amyloids are fibers expressed by bacteria on the cell surface in order to initiate biofilm formation [7].

### Chassis organism

Like its relative, *Bacillus subtilis*, a model organism for gram-positive bacteria, *B. megaterium* is a non-pathogenic, soil-inhabiting, spore-forming bacterium. It has been utilized for industrial purposes for decades due to its capability to secrete large amounts of recombinant proteins [8]. Unlike *B. subtilis*, *B. megaterium* can maintain freely replicating plasmids and has no alkaline protease activity [8]. Its ability to maintain plasmids allows us to more rapidly screen through different genetic constructs, and the lack of alkaline protease activity makes it ideal for protein production. In future applications, where bioremediation functions may be coupled to detection, the ability to secrete proteins in large quantities may be beneficial.

*B. megaterium* forms biofilms naturally, but in order to have enhanced cell-cell or cell-surface adhesion we first identified amyloid proteins, known to promote biofilm stability [7,9,10], that will express well in *B. megaterium*. The first amyloid protein we wanted to test is the functional *E. coli* amyloid curli that is made up of the major subunit CsgA and the minor subunit CsgB [7]. Curli are well-studied proteins that, when exported, form a net-like structure that becomes the major component of the biofilm matrix [7]. The second amyloid protein we wanted to test was TasA, from the gram-positive bacterium *B. subtilis* [9]. It is not known how much amyloid protein has to be expressed in order to form cell aggregates. An inducible system is a straightforward way to tune the protein expression, by simply titrating the inducer, and thereby evaluate how much induction is appropriate.

## Results

### Xylose-inducible constructs were transformed into B. megaterium

After assembling and confirming the validity of the constructs by Sanger sequencing, they were transformed into the *B. megaterium* ATCC12872 as described in the Materials and Methods section. After transformation, the plasmid was purified from *B. megaterium* cells and sequenced again. Six out of the seven proposed constructs were successfully transformed into *B. megaterium* (see Supplemental data A).

### Western blot reveals localization of CsgA and TasA

In order to evaluate protein expression and localization, the His-tagged amyloids were separated on a polyacrylamide gel, transferred to a nitrocellulose membrane, and identified by immunostaining. Based on amino acid sequence, the calculated molecular weight of CsgA is 17.89 kDa and the calculated molecular weight of CsgA + linker is 18.47 kDa. For both CsgA-constructs, only the cell pellet fraction contained any protein, suggesting that CsgA is not being exported (Fig. 1A, lanes 1-4). TasA + linker has a calculated molecular weight of 31.77 kDa. Based on immunostaining, TasA is present both in the supernatant and the pellet, suggesting that the protein is being exported (Fig. 1A lane 5 and 6). The band below the pellet fraction TasA (Fig 1A, lane 6) could be a degradation product of TasA.

**Figure 1.**
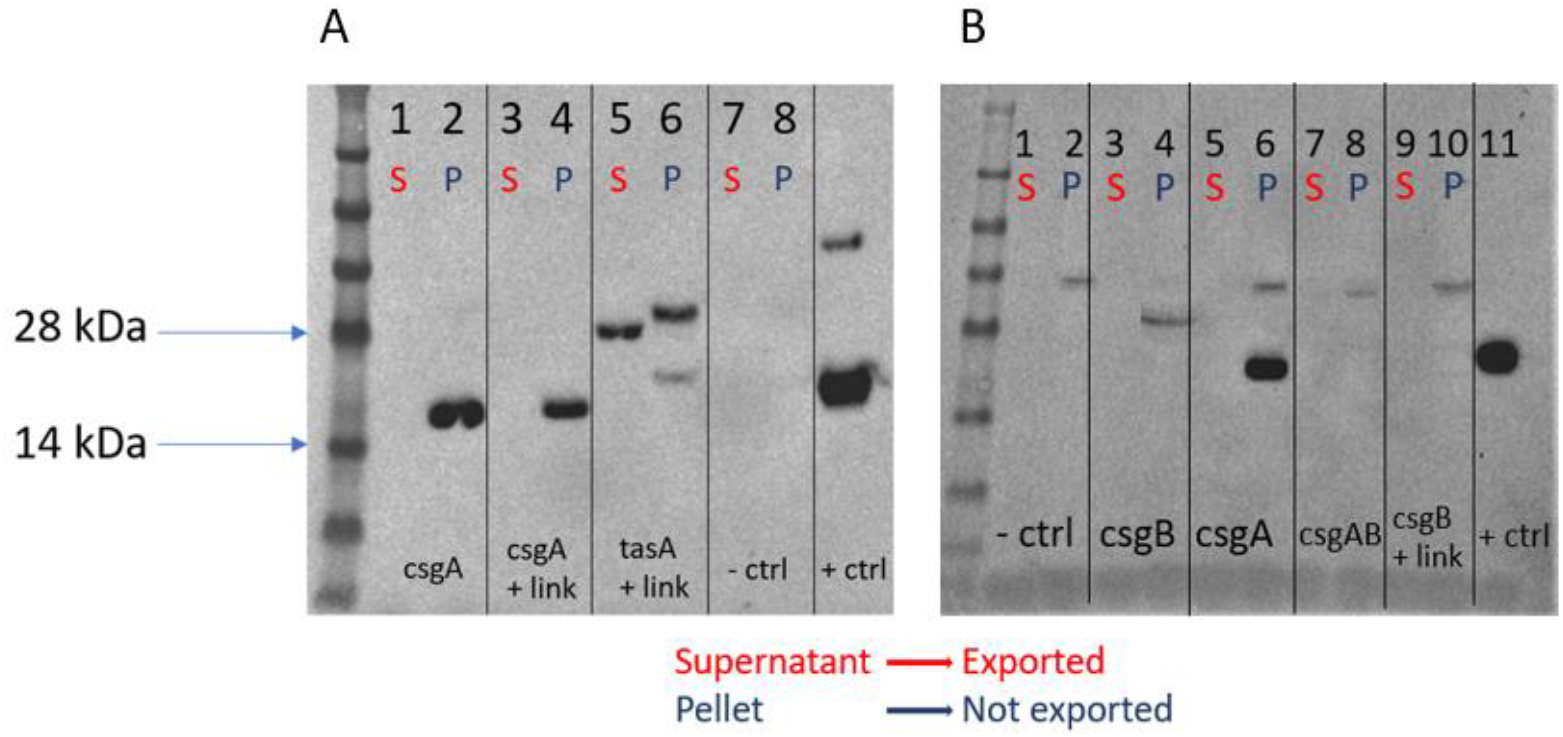
(A) Western blot for supernatant and cells of csgA- and tasA-containing constructs. (B) Western blot for supernatant and cells of csgA, csgB, csgB + linker and csgAB constructs.

In order to confirm that the size difference between TasA in the supernatant and in the cell pellet could be explained by cleavage of the export tag, the SDS-PAGE gels were calibrated using the ladder proteins of known size. The migration distances of the proteins were measured and then plotted against their known molecular weights and the plot was fit to a linear regression. The molecular weight of the tag is 2.89 kDa. The calculated difference between the internal protein and exported protein, using the calibration, is 2.74 kDa, which suggests that the export tag is likely cleaved.

Together these results show that both TasA and CsgA are expressed but only TasA is fully exported.

### Western blots suggest csgB and csgAB are not expressed

In contrast to CsgA and TasA, we found that CsgB and CsgAB were not expressed by *B. megaterium* cells. Based on amino acid sequence, the calculated molecular weight of CsgB is 18.77 kDa, the calculated molecular weight of CsgB + linker is 19.35 kDa and the calculated molecular weight of CsgAB is 36.66 kDa. Based on immunostaining, none of these gene products were present in the cell pellet (Fig. 1B). This suggests that neither CsgB nor CsgAB is expressed. Bands of approximately 38 kDa are visible from the pellet of the negative control on the Western blot, CsgB + linker, CsgA and CsgAB (Fig. 1B lane 2, 6, 8 and 10). These bands are likely an artifact of nonspecific binding or over-developing the membrane. The band crossing lane 4 is a technical artifact caused by a piece of dried polyacrylamide gel on the membrane.

### Western blot reveals tunable expression of TasA

In order to determine if different induction concentrations yielded different concentrations of exported amyloid, samples were taken for induced TasA-expressing cells at 0, 3, 10 and 33.3 mM xylose. Based on immunostaining, the expression of TasA is increased with induction and no protein was expressed from the TasA-expressing cells to which no xylose was added (Fig. 2A lane 1 and 2) which means that the leak of the circuit is low. Based on the band intensities we estimated (see Materials and Methods for details) relative exported protein (Fig 2B, left) and the fraction of exported protein (Fig 2B, right) and found that the fraction of exported protein decreases with induction.

**Figure 2.**
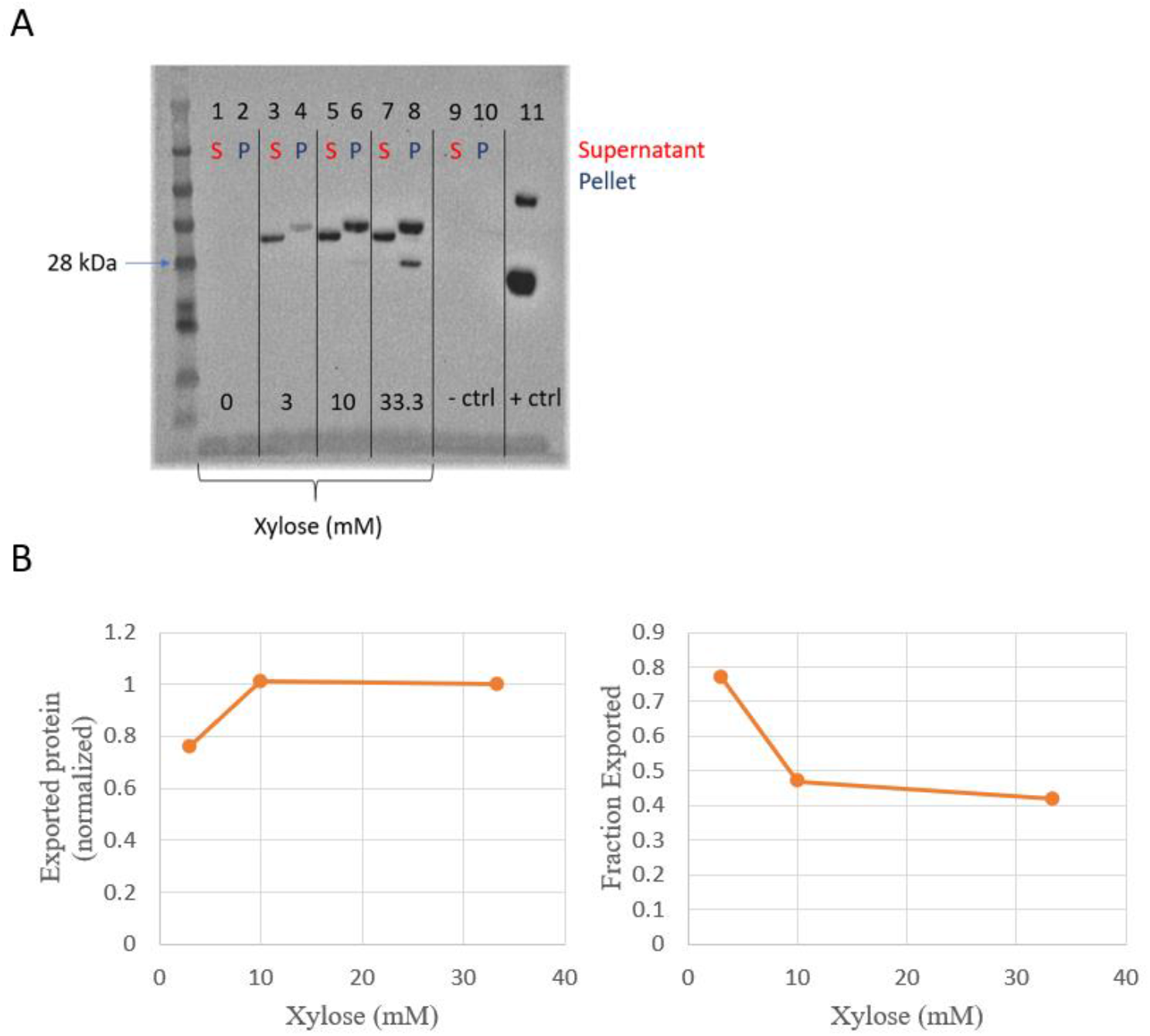
(A) Western blot for supernatant and pellet of tasA-containing cells at different induction concentrations. (B) Relative exported protein (left) and fraction of exported protein (right).

### Flask cultivation endpoint OD600 measurements suggest csgB is being expressed

Despite the fact that CsgB was not present on the Western blot, we observed that cells transformed with the *csgB* construct displayed a lower optical density (OD600) reading 3 hours after induction with 33.3 mM xylose compared to an uninduced condition. This behavior is consistent with the behavior of cells containing *tasA* and *csgA* constructs, but not with cells containing an empty control vector or the *csgAB* construct (Table 1). This result suggests that CsgB, despite it not appearing in the Western blot (Fig. 1B), might still be causing expression burden on the cells and yields further evidence that *csgAB* is not being expressed in the cells. Some possibilities that may explain this penalty are protein degradation, cleavage of the His-tag, or the induction level being too high [11]. Furthermore, the samples were probed for expression at 3 hours after induction. This corresponds to growth curve data where cell density is low.

**Table 1.** Final measured OD600 for construct-containing cells grown in flasks.

| Construct | Final OD600 (induced) | Final OD600 (uninduced) |
| --- | --- | --- |
| *csgA* | 0.816 | 1.72 |
| *csgB* | 0.852 | 2.14 |
| *tasA* | 1.24 | 1.71 |
| *csgAB* | 0.598 | 0.634 |
| pMM1522 empty vector | 1.96 | 1.97 |

### Plate reader induction experiments reveal growth burden caused by expression of CsgA and CsgB, but not TasA

In order to evaluate the growth burden inferred by induction we grew our construct cells at different xylose concentrations. *tasA*-containing cells were grown in xylose concentrations ranging from 0-33.3 mM. *csgA*-, *csgB*- and pMM1522-containing cells were grown in beef extract containing xylose concentrations over a range of 0-25 mM. All *tasA*-construct growth curves, irrespective of xylose concentration, look similar, suggesting little to no growth penalty when expressing the amyloid protein (Fig. 3B). This also applies to the empty vector cells (Fig. 3D). The *csgA*-construct and *csgB*-construct growth curves show higher burden for higher induction concentrations (Fig. 3A and 3C). In addition to this, the growth curves show a recovery at approximately 8 hours of growth. This is not due to mutational breakage of the circuit, but rather due to cellular consumption of the xylose in the growth medium (see Supplemental data D).

**Figure 3.**
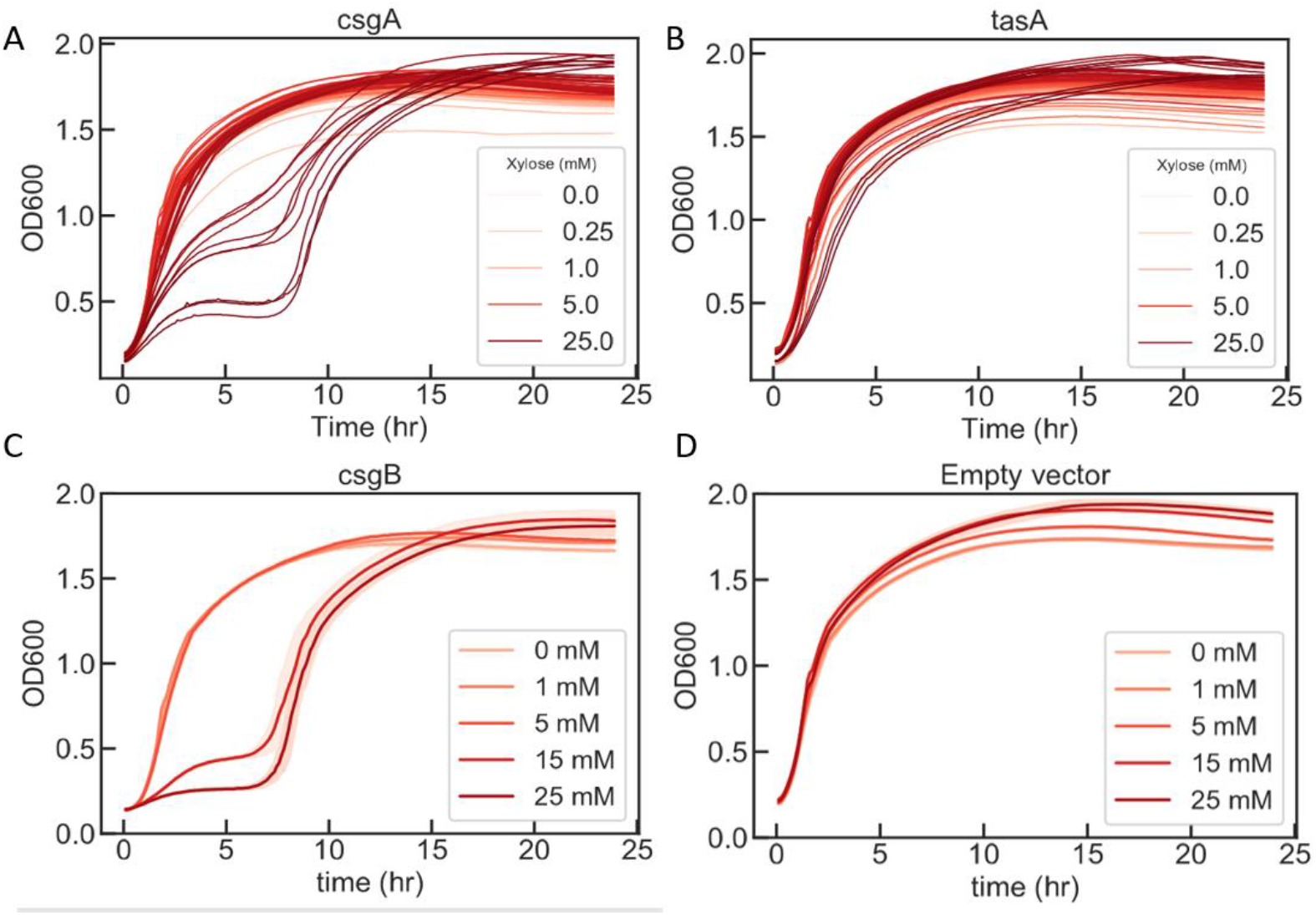
OD600 measurements of construct cells grown in beef extract in different xylose concentrations for 24 hours. (A) csgA-containing cells. (B) tasA-containing cells. (C) csgB-containing cells. (D) pMM1522 empty vector-containing cells.

To further evaluate the growth burden associated with producing amyloid proteins the growth curves from the plate reader induction experiments were fit to a four-parameter logistic regression (Fig. 4A), as described in Materials and Methods. When the estimated growth rates and carrying capacities for the *tasA*-containing cells are plotted against the xylose induction concentrations, there is only a small decline for the highest induction concentrations and that the carrying capacity is similar for all induction concentrations (Fig. 4C). Since the biphasic curve shape for the *csgA*-containing cells made it difficult to fit the data to a logistic growth curve, we chose to take a single data point at 8 hours before the second growth phase. The OD600 at that time point was plotted against the xylose induction concentration for *csgA*- and *tasA*-containing cells grown in LB (Fig. 4B). We observe that the OD600 decreases with increasing xylose concentration for *csgA*-containing cells.

**Figure 4.**
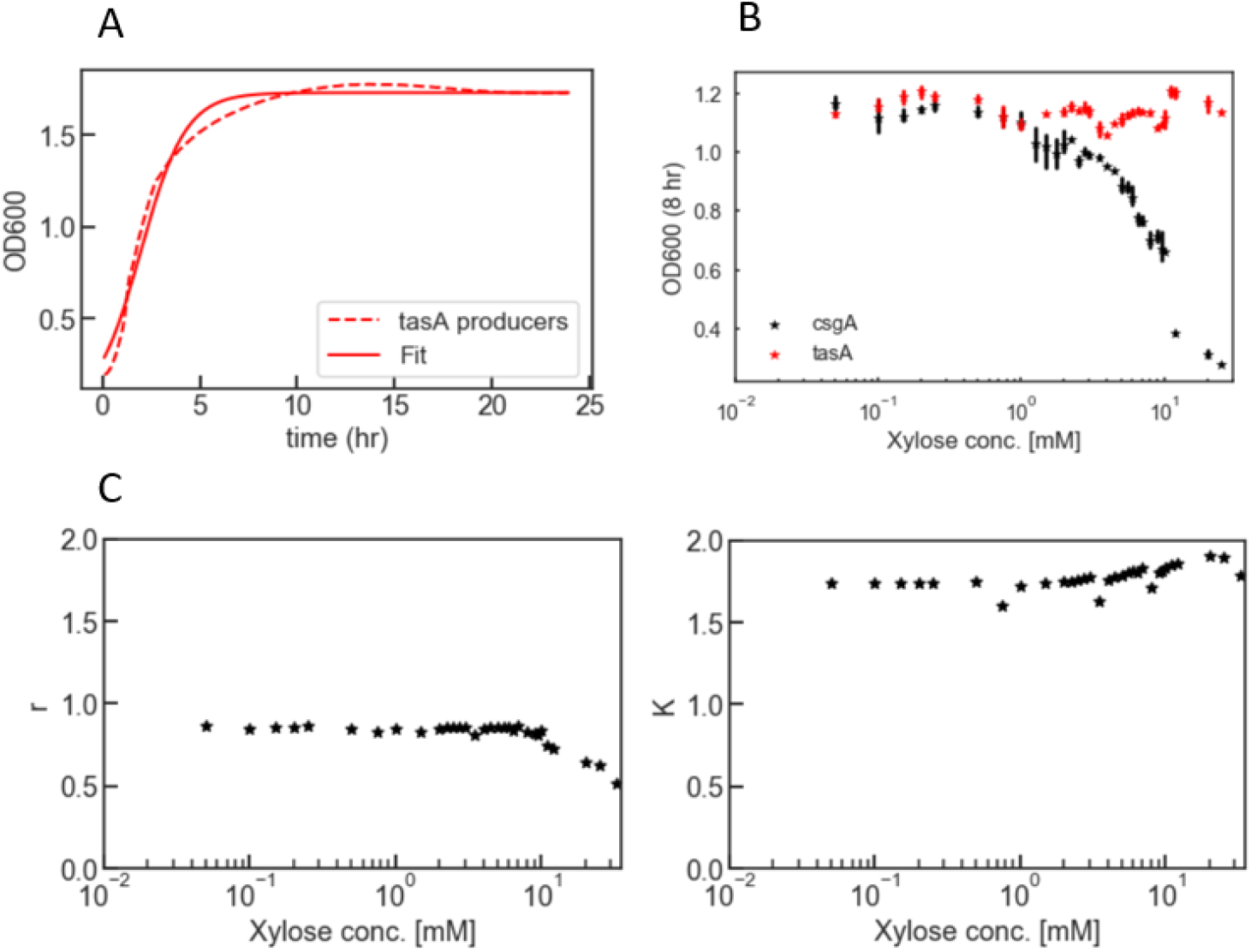
Curve fits for csgA and tasA constructs. (A) Example of logistic curve fit for tasA-containing cells. The data from one induction concentration (20 mM) is fitted to Equation 1. (B) Measured OD600 at 8 hours for csgA- and tasA-containing cells grown in LB. (C) Growth rate against xylose concentration and carrying capacity against xylose concentration for tasA-containing cells.

### Initial nickel bead column experiment shows minimal adhesion effect of TasA expression

To evaluate if the expression of TasA increased cell-cell adhesion, cells expressing His-tagged TasA were evaluated for their ability to bind nickel resin (see details in Methods subsection “Nickel bead column assay”). The protocol is outlined in Fig. 5. If the amyloids form a net-like structure around the cells, binding of the hexahistidine-tag to the nickel resin should also retain some of the cells. After counting the colonies, the ratio between cells from the resin portion and the total number of cells (cells in flow-through and from the resin) was calculated. A high ratio means that the amyloid is promoting cell adhesion to the beads. As expected, the ratio is higher in the induced condition, however although the effect was statistically significant, the magnitude of the effect size is very small (see Fig. 5B and Supplemental data E). This suggests that cell-cell adhesion, as measured by the nickel binding assay, is indeed promoted by expression of TasA but not to a large extent.

**Figure 5.**
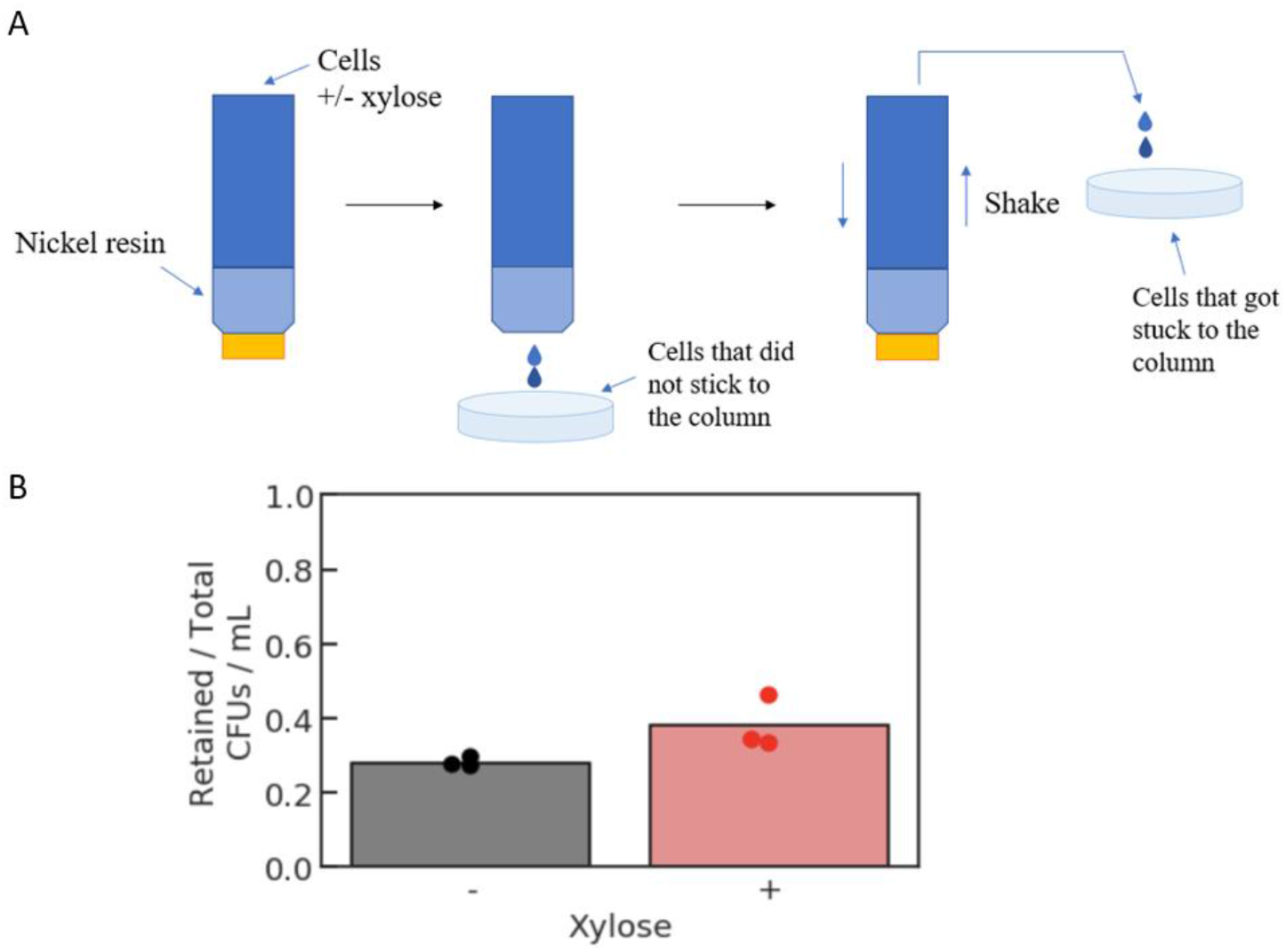
(A) Overview of Nickel bead column assay. (B) Calculated ratios for induced and uninduced cells from three experiments.

## Discussion

The effect from cell-cell adhesion by expression of TasA is very small but could potentially be larger if the assay protocol was optimized. The samples for the Western blots were taken from cultures that had been growing at 37°C with shaking. Ideally the experiment conditions should be more similar to these conditions. One adjustment could be to let the cells grow for a few hours after induction before loading them to the column. This would allow the cells to produce and export more protein before encountering the beads. Another adjustment could be to vary the volume of the beads to prevent the column from being “overloaded” with cells. In addition to improving the nickel bead column assay, alternative assays should be identified to check for adhesion induced by TasA. One option is to develop an assay using microscopy. If improved cell-cell adhesion can be confirmed by other assays than the current one, the next step is to confirm inducible biofilm formation. And lastly, robustness in environmental contexts with and without biofilm should be tested in order to determine the benefit of biofilm formation to survivability.

The construction of the *tasA*-circuit means that the goal of creating an inducible amyloid expression gene in *megaterium* was satisfied. Furthermore, the fact that TasA is being exported (Fig. 1A) and shows minimal growth burden at high expression (Fig. 3B) suggests that TasA is a promising choice for the continued development of an inducible cell-cell adhesion circuit for *B. megaterium*. In contrast to TasA, CsgA could not be successfully expressed and exported. Curli are normally expressed in *E. coli*, which require a complicated machinery for export. This may explain why we do not observe secretion of the curli protein in *B. megaterium*, despite the secretion tag. This suggests that the expression of exogenous genes from closely related species is preferable.

Using synthetic biology to engineer systems for environmental sensing and remediation is of great importance. A step on the way towards building these systems is finding ways to increase the robustness of microbial communities in harsh environments. In this study, we built a xylose inducible amyloid expression circuit for *B. megaterium* with the aim of promoting cell-cell adhesion as a first step in the direction towards this goal.

## Materials and Methods

### Circuit design

The xylose-inducible genetic construct was designed using the plasmid pMM1522, which has a xylose inducible promoter (Fig. 6 and Supplemental data A). The gene sequences for each amyloid protein (CsgA, CsgB, CsgAB and TasA) were codon optimized and a *lipA* export tag and linker sequence, optimized for *B. megaterium* [12], was added to the N-terminus in order to export and cleave the tag. A hexahistidine tag was added at the C-terminus in order to evaluate protein expression by Western blotting.

**Figure 6.**
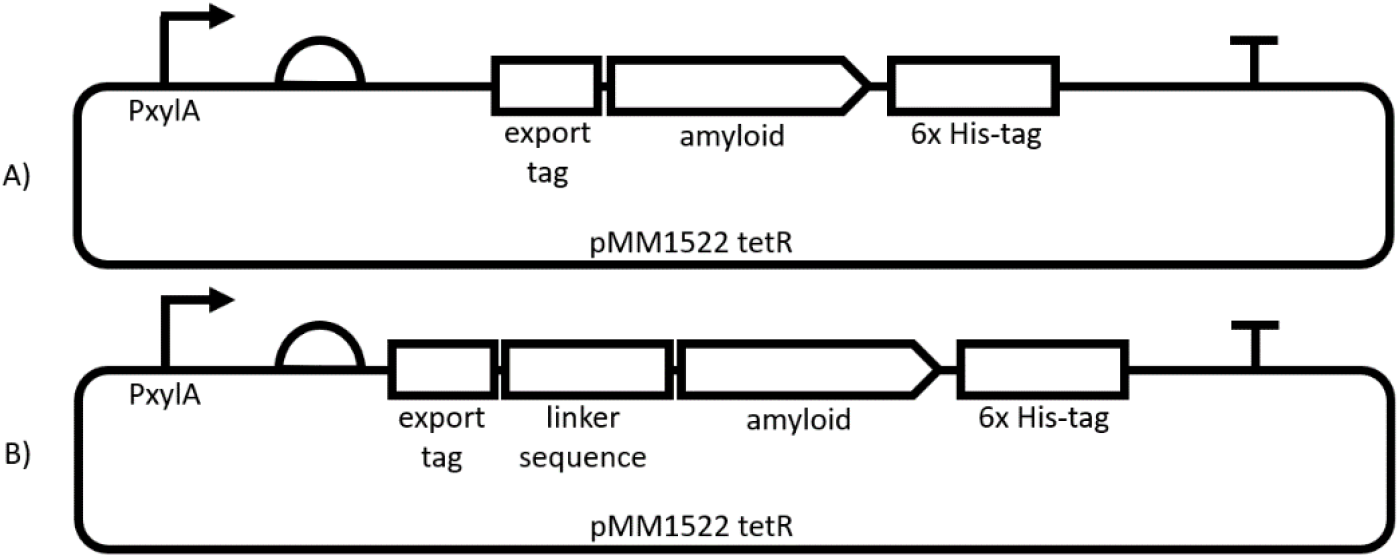
Schematic figure of construct design. (A) Each construct contains a xylose inducible promoter, a strong expression ribosome binding sequence (RBS), an export tag, the amyloid gene, a hexahistidine-tag and a terminator. (B) For some of the constructs, the export tag is followed by a linker sequence.

### UNS-cloning and transformation of *E. coli* cells

The parts described in Figure 1 were assembled using homologous unique nucleotide sequences (UNS) and Gibson assembly [13,14]. DNA sequences containing an RBS, a secretion tag, a linker sequence, the gene of interest and a His-tag were synthesized by idtDNA. UNS sequences were added to the 3’ and 5’ ends of each part by PCR-amplification. The PCR products were separated on a 1% agarose gel and purified using a QIAquick Gel Extraction Kit (Qiagen). The parts were assembled using the NEB HiFi DNA Assembly kit (New England Biolabs) with an incubation time of one hour at 50°C. A volume of 1 μL from each assembly reaction was transformed into *E. coli* chemically competent cells and plated onto LB-agar containing carbenicillin (100 μg/ml) and grown overnight at 37°C. The correct assembly of each construct was confirmed by Sanger Sequencing.

### Transformation of circuit into *B. megaterium*

#### Preparation of *B. megaterium* protoplasts

Cultures of *B. megaterium* cells were grown overnight in LB. A volume of 1 ml of the overnight culture was used to inoculate 50 ml of LB in a baffled flask. The cells were incubated at 37°C at 250 rpm shaking until OD600 reached ~1. The cells were pelleted by centrifugation at 4°C for 15 minutes and 3000 rcf and resuspended in 5 ml freshly prepared SMMP containing 2 mg lysozyme. The cells were incubated at 37°C for 30 min with smooth shaking. The protoplasts were then harvested by centrifugation at 1300 x g for 10 minutes. The supernatant was decanted, and the protoplasts were suspended in 5 ml of SMMP. The cells were washed once more and then suspended in 4 ml SMMP and 1.25 ml of 50 % glycerol was added. The cells were either used directly or stored at −80°C.

#### Transformation of *B. megaterium* protoplasts

500 μl of protoplasts was mixed with 5 μl of DNA in 15 ml tubes. 1.5 ml of PEG-P was added and mixed carefully by inverting the tube. The mixture was incubated for 2 minutes at room temperature. 5 ml of SMMP was added and mixed by carefully inverting the tube. The cells were then harvested by centrifugation at 1300 rcf for 10 minutes. The cells were resuspended in 500 μl SMMP and incubated at 30°C for 45 minutes without shaking followed by 45 minutes with smooth shaking. The cells were then mixed with 2.5 ml CR5 top agar and poured onto plates with selection and grown overnight at 30°C.

### Evaluation of amyloid expression

#### Xylose induction plate reader experiment

Starting from an individual colony, 5 ml of LB containing 10 ug/mL of tetracycline was inoculated and grown overnight. The overnight culture was then diluted 1:100 into fresh 5% beef extract medium (see Supplemental data C for explanation of choice of growth medium) and antibiotic and grown up to mid-log (OD600 0.4-0.7). The cells were then diluted 1:2 into a 96 well with final xylose concentrations ranging from 0 to 33.3 mM. A Biotek Synergy H1 plate reader was used with a 96 well matriplate (DOT Scientific MGB096-1-2-LG-L) at 37°C with maximal linear shaking and OD600 measurements were taken every 7 minutes for 24 hours.

#### Western blot

Cells were grown overnight from a glycerol stock. The cells were then diluted 1:100 into LB containing 20 μg/ml tetracycline. The cells were split into two tubes and xylose to a final concentration of 33.3 mM was added to one of the tubes. Water was added to the second tube to the same final volume. The cells were grown at 37°C for 3 hours and the final OD was measured. The cells were pelleted, and the supernatant was collected. The cells were washed with PBS once and then resuspended in PBS containing 1 mg/ml lysozyme and incubated at 37°C for 10 minutes. The samples were frozen at −80C for later use.

Samples of 30 μl each were thawed and mixed with 10 μl of 4x NuPAGE LDS sample buffer (ThermoFisher). The samples were incubated at 98°C for 10 minutes and 10 μl of each mixture and a SeeBlue Plus2 pre-stained protein ladder (Thermofisher) were loaded onto a NuPAGE 4-12 % Bis-Tris polyacrylamide gel (ThermoFisher). Proteins were separated by electrophoresis for 30 minutes at 200 V. The proteins were transferred from the gel to a nitrocellulose membrane using a iBlot 2 device (ThermoFisher) and the membrane was immunostained using an iBind instrument (ThermoFisher). The primary antibody was Rabbit anti-hexahistidine tag and the secondary antibody was Goat anti-rabbit conjugated to horseradish peroxidase. The blot was developed for 1-2 minutes using a SuperSignal West Pico PLUS Chemiluminescent substrate (ThermoFisher) and imaged.

Band intensities were measured using ImageJ [15] in order to calculate an estimate of the relative amounts of exported protein. The relative exported protein was calculated by normalizing the band intensities by the band intensity of the 33.3 mM induction level. The fraction of exported protein was calculated by dividing the intensity of the supernatant band with the intensity of the supernatant band and the intensity of the pellet band after subtracting background.

#### Curve fitting

The growth curves from the plate reader induction experiment were fit to a logistic curve (Equation 1) using Python.

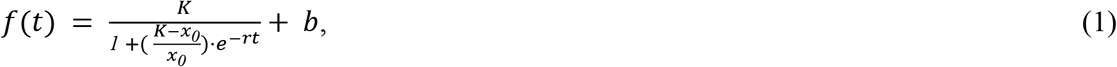

where *K* is the carrying capacity, *r* is the growth rate, *x*_*0*_ is the initial cell concentration, *t* is the time and *b* is the floor term (accounting for background). Growth rates and carrying capacities for different induction concentrations were estimated from the curve fit (using Python).

#### Nickel bead column assay

Starting from an individual colony, an overnight culture was grown in LB. The cells were diluted 1:100 in 5% beef extract and grown to mid-log. A column was loaded with 1 ml nickel resin and washed with 3 ml of water and then 10 ml of 5% beef extract. 10 ml of cells were carefully layered on top of the resin and incubated for 4 hours at room temperature with or without xylose inducer (33.3 mM). After four hours, the column was unplugged and the flow-through was collected. After that, the column was plugged and filled with 10 ml beef extract and the resin was mixed with the extract. The flow-through portion and the nickel portion were then plated onto separate tetracycline plates in different dilutions and grown overnight at 37°C. The next day, the colonies of the plates with an appropriate number of colonies (~30-200), were counted. The ratio between the number of colonies on the plate with the resin portion and the total number of cells (resin portion and flow-through portion) was then calculated. The ratios for the induced and uninduced case were then compared.

#### Statistical analysis of nickel bead column assay data

CFU/mL values for retained cells and flow-through cells were used to compute ratios of retained cell density / total cell density for each replicate in each condition. The ratios in the induced condition were assessed for statistical significance against the ratios in the uninduced condition by using a nonparametric permutation test where the null hypothesis is that the distribution of ratio values in the induced and uninduced condition are equal. The difference of the means in the two conditions was used as the test statistic The test was performed as follows: we pooled the ratio values for both conditions together, shuffled the data, and randomly labeled half of these values as “induced” and the other half as “uninduced”, and calculated the difference between the mean of the “induced” set and the mean of the “uninduced” set. This process was repeated 100,000 times. The p-value is the frequency of the event that the difference of means computed after shuffling was greater than or equal to the difference of means observed empirically in our experiment. The test was implemented using a custom Python script.

## Acknowledgments

This research is supported by the Institute for Collaborative Biotechnologies through contract W911NF-19-D-0001 from the U.S. Army Research Office. The content of this report does not necessarily reflect the position or the policy of the Government, and no official endorsement should be inferred.

