## Supplemental data for "Construction of an inducible amyloid expression circuit in *Bacillus megaterium:* A case study with CsgA and TasA"

### A. pMM1522 Plasmid map and xylose-inducible adhesion constructs

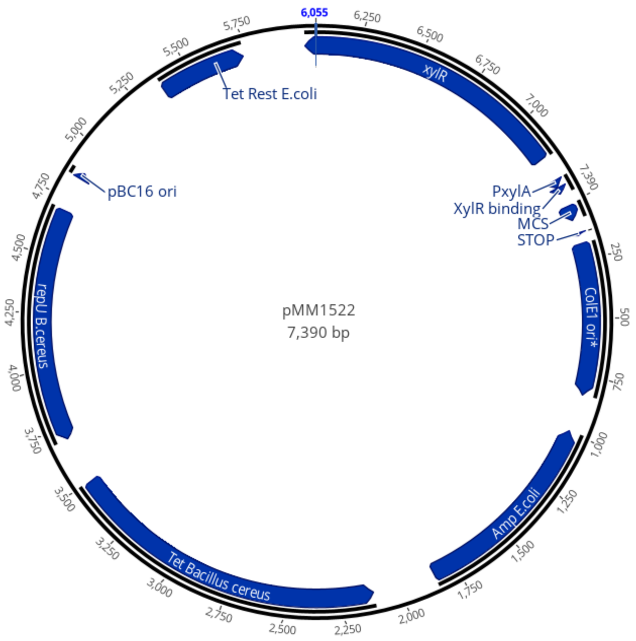


**Table A**. Xylose- Inducible Adhesion Constructs.

| **Construct name** | **Description** | **Transformed into *E. coli*** | **Transformed into *B. megaterium*** |
|---|---|---|---|
| *csgA* | Major curli subunit | ✓ | ✓ |
| *csgA* + linker | *csgA* and linker sequence | ✓ | ✓ |
| *csgB* | Minor curli subunit | ✓ | ✓ |
| *csgB* + linker | *csgB* and linker sequence | ✓ | ✓ |
| *csgAB* | *csgA* followed by *csgB* | ✓ | ✓ |
| *tasA* | *B. subtilis* amyloid | ✓ | ✗ |
| *tasA* + linker | *tasA* and linker sequence | ✓ | ✓ |

### B. Gel image of parts for Gibson assembly

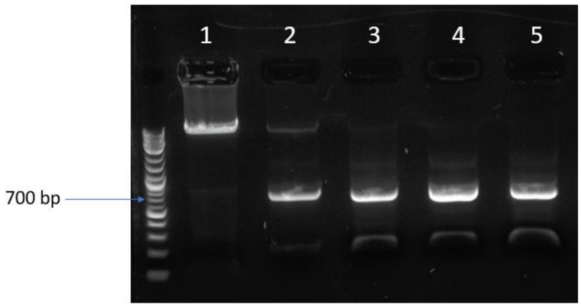


**Figure B1. Gel image of parts for Gibson assembly.** Picture of agarose gel bands for (1) the backbone piece, (2) csgA, (3) csgB, (4) csgA + linker, (5) csgB + linker.

### C. Media optimization

***Plate reader induction experiment and visual phenotype reveal B. megaterium forms clusters when grown in LB***

In order to further investigate the relationship between amyloid protein expression and cell growth dynamics, we measured growth over time at different induction levels for each of our constructs. The growth curves for cells grown in LB showed a high degree of variation in OD600 between 2 and 7 hours (Fig. 9). We surmise that this variation may be due to cell clumping. We observed visual clumping for cells grown in 5 mL of LB, at mid-log phase (3 hours). These cultures returned to homogeneity at stationary phase. These observations mirror those recorded in the plate reader. We could not find anything in the literature that explains why this is happening.

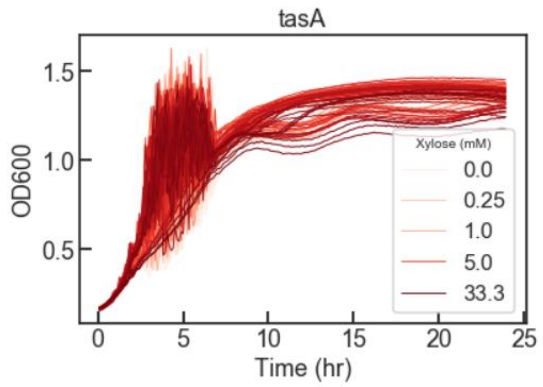


***Figure C1. OD600 measurements of tasA-containing cells grown in LB in different xylose concentrations***. *tasA-containing cells were grown in LB for 24 hours in xylose concentrations from 0 to 33.3 mM in triplicates. All concentrations and triplicates are plotted. Around 3-5 hours, the OD600 measurements are very noisy for all induction concentrations due to cell clumping in the logarithmic growth phase.*

### D. Follow-up experiments on plate reader induction experiment

***Growing cells in the supernatant suggests xylose is being consumed after 7 hours of growth***

After excluding the possibility that the circuit had incurred mutations, we reasoned the biphasic nature of the growth curves may be due to metabolism of the xylose, thereby repressing expression of the csg proteins.  To test this, *csgA*- and *csgB*-containing cells were grown in beef extract containing xylose concentrations ranging from 0-33.3 mM. One row in the plate was filled with 300 µl beef extract (33.3 mM xylose) without any cells to make sure heating of the medium did not affected the xylose. The cells were grown at 37ºC for 22 hours and then centrifuged to separate the cells from the supernatant. The supernatant, including the media without added cells, was then transferred to a new plate and fresh cells, grown to mid-log, were added to the plate and grown for 24 hours. The fresh cells that grow in the supernatant from cells grown in 33.3 mM xylose do not show biphasic growth behavior, while cells grown in the 33.3 mM xylose medium have the same recovery at 7 hours as the cells in the first plate. This suggests that the xylose is completely consumed at 7 hours when the cells can start to grow normally again.

***Sequencing results reveal the csgA-circuit is intact after 24 hours of growth***

To determine if the biphasic growth curve shape (Fig. 4A and 4C) could be explained by a large deletion or insertions in the genetic circuit after 7-8 hours, the *csg* gene was PCR-amplified from construct containing cells. The PCR product was separated on an agarose gel. Bands of the expected molecular weight were present for all xylose concentrations (Fig. D1 and D2). The PCR product was then sequenced to determine if any point mutations had occurred. The sequencing results showed that the construct is unchanged after the construct-containing cells have been grown for 24 hours.

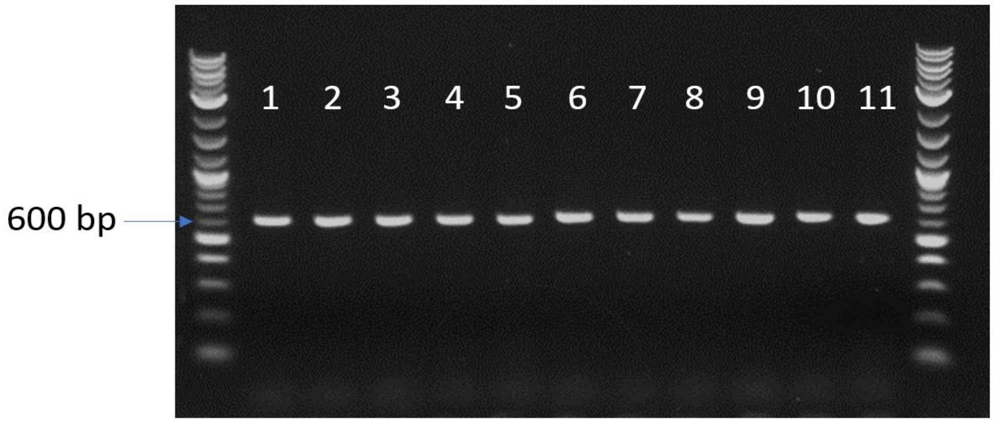


***Figure D1. csgA plate reader PCR.*** *(1) 0 mM xylose in LB, (2) 2 mM xylose in LB, (3) 6 mM xylose in LB, (4) 20 mM xylose in LB, (5) 25 mM xylose in LB, (6) 0 mM xylose in beef extract, (7) 2 mM xylose in beef extract, (8) 6 mM xylose in beef extract, (9) 20 mM xylose in beef extract, (10) 25 mM xylose in beef extract and (11) individual csgA-containing cell colony from agar plate. The expected length of csgA is 654 base pairs.*

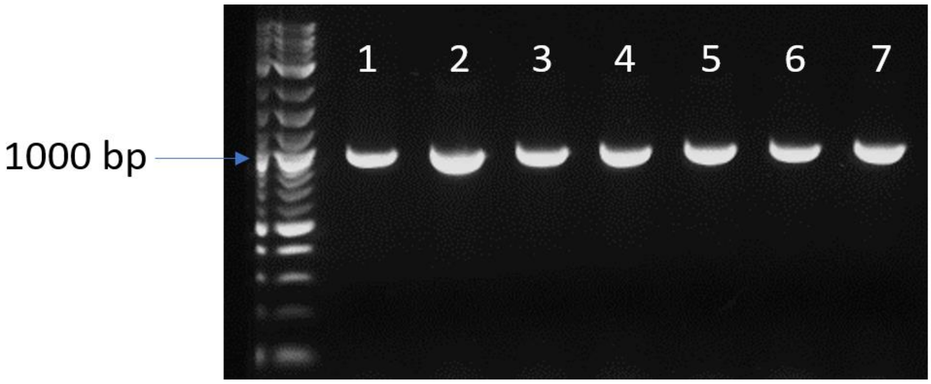


***Figure D2. tasA plate reader PCR.*** *(1) 0 mM xylose in beef extract, (2) 2 mM xylose in beef extract, (3) 6 mM xylose in beef extract, (4) 20 mM xylose in beef extract, (5) 25 mM xylose in beef extract, (6) 33.3 mM xylose in beef extract and (7) individual tasA-containing cell colony from an agar plate. The expected length of tasA is 1002 base pairs.*

**E. Nickel bead column assay - data**

**Table E.** Ratios (Retained cells/All cells) for induced and uninduced cells in the Nickel column assay and calculated p-value from permutation test.

| **Replicate** | **Uninduced** | **Induced** |
|---|---|---|
| 1 | 0.296 | 0.333 |
| 2 | 0.272 | 0.462 |
| 3 | 0.276 | 0.3425 |
| Mean | 0.282 | 0.379 |
| p-value | 0.01655 | |

**F. DNA sequences**

| **Name** | **Description** | **Sequence** |
|---|---|---|
| UNS3F_seq | Forward primer for sequencing of constructs | CGCACTGGAAACATC |
| UNS3R_seq | Reverse primer for sequencing of constructs | GTTTGTTTCCGTCTACG |
| UNS3_FOR | Forward primer for adding UNS3 to 5'-end of construct | GCACTGAAGGTCCTCAATCGCACTGGAAACATCAAGGTCGTAGAGTATAAGGAGGTTACT |
| UNS2_REV | Reverse primer for adding UNS2 to 3'-end of construct | GCTTGGATTCTGCGTTTGTTTCCGTCTACGAACTCCCAGCTTAATGGTGATGGTGATGGT |
| UNS4_FOR | Forward primer for adding UNS4 to 5'-end of construct | CTGACCTCCTGCCAGCAATAGTAAGACAACACGCAAAGTCTAGAGTATAAGGAGGTTACT |
| UNS4_REV | Reverse primer for adding UNS4 to 3'-end of construct | GACTTTGCGTGTTGTCTTACTATTGCTGGCAGGAGGTCAGTTAATGGTGATGGTGATGGT |
| UNS2_pMM1522_FOR | Forward primer for adding UNS2 to the linear plasmid backbone | GCTGGGAGTTCGTAGACGGAAACAAACGCAGAATCCAAGCACCTCGCTAACGGATTCACC |
| UNS3_pMM1522_REV | Reverse primer for adding | CGACCTTGATGTTTCCAGT |

| | | |
|---|---|---|
| | UNS3 to the linear plasmid backbone | GCGATTGAGGACCTTCAGT GCATTGAGTTAGTTTGTTT ATC |
| SPlipA export tag | Export tag sequence | ATGGCCAATCAACCGACGA AATACCGCAAGTTTGTTGT AGGAGCGGCGAGCGCGGC CTTAGTGGCTTCAGCGGTC GCGCCTGTCGCGTTT |
| SPlipA linker sequence | Linker sequence | GAATCAGTACATAAT |

### G. Media

**2x AB3 (1 L)**
Autoclave for 15 minutes

- 3 g Beef Extract
- 3 g Yeast Extract
- 10 g peptone
- 1 g glucose
- 7 g NaCl
- 7.36 g $K_2HPO_4$
- 2.64 g $KH_2PO_4$

**2x SMM**
pH adjust to pH 6.5 and filter sterilize

- 1 M Sucrose
- 40 mM maleic acid
- 40 mM $MgCl_2$

**PEG-P**
Autoclave for 15 minutes:

- 40% PEG 6000 in 1x SMM

**SMMP**
2 x AB3 and 2 x SMM; mixed 1:1

**Solution A**
pH adjust to pH 7.3 and filter sterilize

- 602 mM
- 58 mM MOPS
- 30 mM NaOH

**Solution B**
Autoclave 15 minutes

- 4 % (w/v) agar agar
- 0.2 % (w/v) casamino acids
- 10 % (w/v) yeast extract

**8 x cR5-salts**

Autoclave

- 11 mM $K_2SO_4$
- 394 mM $MgCl_2$ x 6 $H_2O$
- 3 mM $KH_2PO_4$
- 159 mM CaCl2

**cR5 top agar (2.5 ml)**

- 1.25 ml Solution A
- 713 µl Solution B
- 288 µl cR5-salts
- 125 µl 12 % (w/v) proline
- 125 µl 20 % (w/v) glucose
